# Quiescent Body as a Reversible Subcellular Structure Formed in Non-growing Bacterial Cells

**DOI:** 10.1101/107466

**Authors:** Jiayu Yu, Yang Liu, Huijia Yin, Zengyi Chang

**Affiliations:** The State Key Laboratory of Protein and Plant Gene Research, School of Life Sciences, Peking University, Beijing 100871, P.R. China; Center for Protein Science, Peking University, Beijing 100871, P.R. China

## Abstract

Bacterial cells take a variable lag time and maintain a multi-drug tolerant non-growing state before resuming growth when encountering growth-supportive conditions. Some of them exhibit an extraordinarily long lag, as pathogenic persisters do. It remains unknown on what determines lag time duration. Here, we unveiled a subcellular structure, termed quiescent body, that is formed in bacterial cells entering non-growing state and sequesters selected proteins essential for cell growth. Their formation occurs progressively in each cell and heterogeneously among individual cells. They only dissolve in re-growing cells to release proteins for immediate re-functioning when conditions become fit. Quiescent body, whose degree of formation is highly correlated with duration of lag time or level of antibiotic tolerance, apparently functions as a biological timer for bacterial growth resumption. Further, suppressing their formation, which directly relies on cellular respiration complexes, or promoting their dissolution might be a viable strategy to eradicate persisters.

## Introduction

Multi-antibiotic tolerance refers to the ability of a bacterium to escape the killing effect of multiple antibiotics (Bigger, 1944; Lewis, 2007). Such multi-antibiotic tolerance is likely caused by the bacteria to maintain in a non-growing “drug-indifference” state (Brauner et al., 2016; Tuomanen et al., 1986), as in general, antibiotics preferentially target the metabolically active growing bacteria but show less effect on those metabolically inactive non-growing ones (Horne and Tomasz, 1977). Bacterial cells would enter a non-growing state when their living conditions become unfit, such as the stationary-phase during culturing in the laboratory (R Kolter et al., 1993). Intriguingly, such a non-growing state can be maintained, although transiently rather than permanently, even when the bacterial cells encounter growth-supportive conditions, such as during the lag phase of their culturing (Levin-Reisman et al., 2010), with the bacteria remain tolerance towards multiple antibiotics (Balaban et al., 2004; Fridman et al., 2014).

Interestingly, even for a genetically identical bacterial cell population the duration of the lag time for individual cells exhibits high heterogeneity (Baranyi, 2002). Although this heterogeneity is viewed to provide a bet-hedging strategy for a bacterial species to adapt to the fluctuating and unpredictable environments (Balaban et al., 2004), the biological cause of this variation remain almost totally unknown. Clinically, bacterial cells manifesting extremely long lag time are referred to as persisters. Due to their tolerance towards different types of antibiotics, persisters have been recognized as one of the greatest challenges in treating infectious diseases (Fisher et al., 2017; Lewis, 2010). In biological terms, it remains unusually difficult to learn about the molecular characteristics of such bacterial cells in the lag phase (as the persisters) due to their low metabolic activity. Although the timing of growth resumption, or difference in the duration of lag time among individual bacterial cells appear to be stochastic (Buerger et al., 2012), recent studies revealed that there seems to be a correlation between the timing of growth resumption and the timing of last entry into the non-growing state for bacterial cells (Jõers and Tenson, 2016). However, it remains largely unknown on what cellular components and/or structures actually determine the duration of the lag time or the timing of growth resumption in individual bacterial cells.

Here, we describe a reversible subcellular structure, termed quiescent body, which is formed in non-growing late stationary-phase bacterial cells and sequesters selected essential proteins, but is disassembled when the cells resume growth and to release the stored proteins for immediate re-functioning. We further demonstrated that both the duration of lag time for bacterial recovery and the level of antibiotic tolerance are strongly correlated to the degree of quiescent body formation.

The discovery of the quiescent body was made by us when performing unnatural amino acid-mediated protein photo-crosslinking analysis in living *E. coli* cells to decipher the assembly pattern of the cell division protein FtsZ, a homolog of the eukaryotic tubulin protein (Erickson, 1998). Under *in vitro* conditions, the FtsZ monomers are known to self-associate into fibrous protofilaments (Erickson et al., 1996), which are believed to further assemble, via a not yet defined manner, into the Z-ring structure in the middle of each parent cell for generating the constriction that leads to the production of two daughter cells. By convention, we should have performed such *in vivo* protein photo-crosslinking analysis of FtsZ only with actively dividing log-phase cells because only in which the Z-ring structure would form. Nevertheless, out of curiosity, we did the same analysis also with the non-dividing late stationary-phase cells. Strikingly, we found that a large portion of the FtsZ monomers, although indeed no longer self-assemble into protofilaments, were present in the insoluble pellet fraction of the cell lysates. Subsequently by live-cell imaging analysis, the FtsZ protein in each late stationary-phase cell was revealed to exist as cell-pole granules or quiescent bodies.

This newly discovered quiescent body structure apparently functions as a biological timer for growth resumption of non-growing bacterial cells. Our findings have immediate clinical applications since blocking the formation or promoting the dissolution of quiescent bodies may provide a viable strategy to eradicate the multidrug-tolerant bacterial pathogen persisters.

## Results

### The FtsZ protein exists in the insoluble pellet fraction of the lysate of non-growing late stationary-phase *E. coli* cells

The crystal structure of the FtsZ protofilaments revealed a head-to-tail longitudinal assembly pattern for the FtsZ monomers (Matsui et al., 2012). We attempted to elucidate whether such self-assembly pattern exists in living *E. coli* cells. For this purpose, we performed *in vivo* protein photo-crosslinking analyses by genetically introducing the photoactive unnatural amino acid *p*-benzoyl-L-phenylalanine (pBpa) (Chin et al., 2002; Ryu and Schultz, 2006) at the determined assembly surface of FtsZ, as we have successfully performed on other proteins (Fu et al., 2013; Wang et al., 2016; Zhang et al., 2011). One variant we prepared, by replacing residue K140 (Li et al., 2013) with pBpa, was FtsZ-K140pBpa, which could substitute the wild-type FtsZ in supporting cell division, as shown in Fig. S1A, indicating its capacity to assemble into a functional Z-ring structure in living cells,

To further determine whether residue K140 is indeed involved in mediating the longitudinal assembly of FtsZ monomers in living cells, we also modified the endogenous *ftsZ* gene to generate FtsZ-Avi variant whose Avi tag can be specifically probed with streptavidin. As shown by the blotting results displayed in Fig. 1A, FtsZ dimers can either form between the FtsZ-K140pBpa and FtsZ-Avi monomers (red arrows) or between two FtsZ-K140pBpa monomers (black arrow) were clearly detected in actively dividing log-phase cells (lanes 2 and 6). These results confirm that residue K140 is indeed located at a self-assembling interface in FtsZ in actively dividing cells.

**Figure 1.**
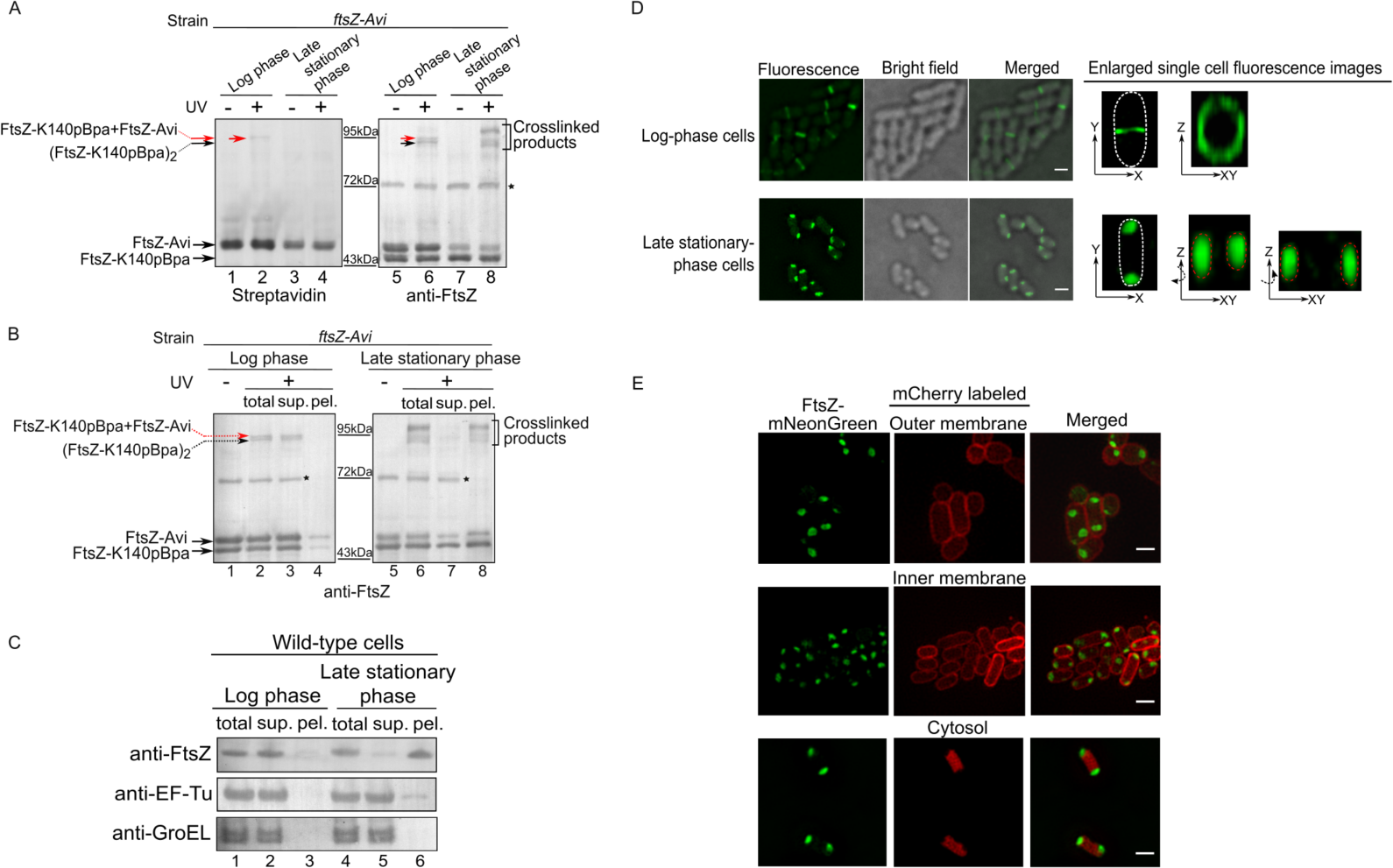
The FtsZ protein in *E. coli* cells exists as a self-assembled oligomer in the actively dividing log-phase but as an insoluble form in the non-growing late stationary-phase. Blotting results for detecting photo-crosslinked products of the FtsZ-K140pBpa variant in log-phase and late stationary-phase *ftsZ*-*Avi* cells exposed to UV light, as probed with streptavidin-alkaline phosphate conjugate or antibodies against FtsZ. The asterisk indicates a non-specific protein band generated when probed with the anti-FtsZ antibodies. Immunoblotting results for detecting the photo-crosslinked products of the FtsZ-K140pBpa variant, as well as the free FtsZ monomers, in the supernatant (sup.) and pellet (pel.) fractions of the log-phase or late stationary-phase *ftsZ*-*Avi* cells, as probed with antibodies against FtsZ. Positions are similarly indicated as in (**A**). Immunoblotting results for detecting the endogenous FtsZ, EF-Tu, or GroEL in the total cell lysate (total), supernatant (sup.), and pellet (pel.) fractions of log-phase or late stationary-phase wild-type *E. coli* cells, as probed with the indicated antibodies. Fluorescence and bright field microscopic images of log-phase (cultured to 6 h; top panel) and late stationary-phase (cultured to 24 h; bottom panel) *E. coli* cells in which the FtsZ-mNeonGreen was heterologously expressed from a plasmid in addition to the endogenously expressed FtsZ. Enlarged single cell fluorescent images are displayed for a better view of the Z-ring (in the log-phase cells) and the cell-pole granules (in late stationary-phase cells). Scale bars, 1 m. Fluorescence microscopic images of late stationary-phase *ftsZ-mNeonGreen* cells whose outer membrane (top), inner membrane (middle), or cytosol (bottom) was separately labeled via the fused mCherry (for the two membranes) or unfused mCherry (for the cytosol). Scale bars, 1m.

As almost none of the previous cellular level studies on the FtsZ assembled Z-ring structure were performed with non-dividing cells, out of curiosity, we also performed the above photo-crosslinking experiments with non-dividing/non-growing late stationary-phase bacterial cells. Surprisingly, the results, also shown in Fig. 1A, revealed there was no photo-crosslinked dimers between FtsZ-K140pBpa and FtsZ-Avi monomers (lane 4). Instead, multiple photo-crosslinked products between FtsZ-K140pBpa and other proteins could be readily detected (lane 8). These results indicate that in such non-dividing/non-growing bacterial cells the FtsZ monomers are prevented to assemble into the protofilaments, likely due to their interaction with other proteins.

In order to identify the proteins interacting with FtsZ in late stationary-phase cells, we then purified the photo-crosslinked products of FtsZ-K140pBpa via immunoprecipitation and subjected them to mass spectrometry analysis. During the purification, we found, strikingly, that almost all the photo-crosslinked products as well as a large portion of the free FtsZ monomers were detected in the insoluble pellet fraction of the late stationary-phase cell lysate (Fig. 1B, lane 8), whereas either the photo-crosslinked FtsZ dimers or the free FtsZ monomers were principally detected in the soluble supernatant fraction of the log-phase cell lysate (Fig. 1B, lanes 3 and 4).

To rule out the possibility that the detection of FtsZ in the pellet fraction was an artifact due to the introduction of pBpa or the Avi tag and/or the UV irradiation, we analyzed the status of the endogenous FtsZ protein in wild-type *E. coli* cells. Our immunoblotting results (Fig. 1C) clearly demonstrated that, similarly, the endogenous FtsZ protein was largely detected in the pellet fraction of late stationary-phase cell lysates (top panel, lane 6). By contrast, as also shown in Fig. 1C, EF-Tu and GroEL were both detected largely in the supernatant fractions (lanes 2 and 5) of either log-phase or late stationary-phase cells. Taken together, these results strongly suggest that the FtsZ protein exists as insoluble forms in the non-growing late stationary-phase *E. coli* cells.

### The FtsZ protein exists as cell pole granules in non-growing late stationary-phase *E. coli* cells

We next attempted to gain further insight into the status of the FtsZ protein in the late stationary-phase cells by performing live-cell fluorescence microscopic imaging analysis. For this purpose, the FtsZ protein fused with the green fluorescent protein mNeonGreen (designated as FtsZ-mNeonGreen) was heterologously expressed in wild-type *E. coli* cells, as previously reported (Ma et al., 1996). We also confirmed that the FtsZ-mNeonGreen was able to effectively incorporate into and, thus, to label the Z-ring structure in the middle of actively dividing log-phase cells (Fig. 1D, top part). We then subjected the non-growing late stationary-phase cells to similar live-cell imaging analysis, and revealed, to our great interest, that the FtsZ-mNeonGreen protein was detected as two granules in each cell, with one at each pole (Fig. 1D, bottom part). By contrast, the similarly expressed unfused mNeonGreen protein itself was evenly distributed in the cytoplasm in either actively dividing log-phase or non-dividing late stationary-phase cells (Fig. S1B). In line with these imaging data, a majority of either the heterologously expressed FtsZ-mNeonGreen or the endogenous FtsZ was detected in the pellet fraction of the late stationary-phase cell lysate (Fig. S1C, lane 6). Collectively, these results demonstrated that the FtsZ protein exists as cell-pole granules rather than as the Z-ring structure in non-growing late stationary-phase bacterial cells.

To further clarify the subcellular localization of the cell-pole granules by dual-color imaging, we first constructed an *E. coli* strain by integrating the *ftsZ*-*mNeonGreen* gene into the genomic rhamnose operon, such that the expression of FtsZ-mNeonGreen was controlled by the rhamnose (Fig. S2A). Similarly, the Z-ring in log-phase cells and the cell-pole granules in late stationary-phase cells could be clearly visualized in the *ftsZ-mNeonGreen* strain when cultured in the presence of rhamnose (Fig. S2B).

We subsequently individually labeled the outer membrane via the OmpA (an outer membrane protein)-fused red fluorescent protein mCherry (Verhoeven et al., 2013) (Fig. 1E, top panel), the inner membrane via the inner membrane anchoring peptide (derived from the nlpA protein)-fused mCherry (Harvey et al., 2004) (Fig. 1E, middle panel), and the cytosol via the unfused mCherry (Fig. 1E, bottom panel) in the *ftsZ-mNeonGreen* cell. These live-cell imaging data clearly showed that each cell-pole granule seems to be rather compact structure that occupied a cytosolic location that was barely accessible to the cytosolic proteins, and that the cell pole granules are closed associated with the inner membrane of the bacterial cells. In line with these observations, we also demonstrated that these cell-pole granules could be isolated as intact entities in the centrifugation pellet of lysed bacterial cells (Fig. S2C).

### The cell-pole granule (or quiescent body) selectively sequesters proteins that are vital for cell growth and division

Insoluble form of proteins was previously reported to accumulate in stationary phase *E. coli* cells (Kwiatkowska et al., 2008; Maisonneuve et al., 2008). Nevertheless, mainly due to the lack of effective ways to examine the status of the proteins present in them, such forms of proteins were naturally assumed to be misfolded. In view of this, we were motivated to find a way to examine whether the FtsZ proteins residing in the cell-pole granules were misfolded or not.

It seemed to us that *in vivo* protein photo-crosslinking analysis as mediated by the unnatural amino acid pBpa would make a feasible approach, in view of its effectiveness for identifying the specific interaction surfaces for two interacting proteins at the amino acid residue resolution (Shiota et al., 2015). Specifically, for a variety of pBpa variants of the FtsZ protein, photo-crosslinked products would be formed with itself or other proteins in surface specific manner if folded properly. To this end, we made use of a pool of pBpa variants of FtsZ that we constructed for unveiling the surfaces on FtsZ monomers for assembling into the Z-ring structure in actively dividing cells.

The results, as shown in Fig. 2A, demonstrated that the pBpa variants of the FtsZ protein apparently interact with other proteins in a surface-specific fashion in non-growing *E. coli* cells. Specifically, when pBpa was placed at residue positions occupying spatial positions adjacent to each other according to the determined crystal structure of FtsZ (Löwe and Amos, 1998), similar patterns of photo-crosslinked products were displayed (e.g., residues 140, 151, 166 and 174, or 31, 47, 51 and 54).

**Figure 2.**
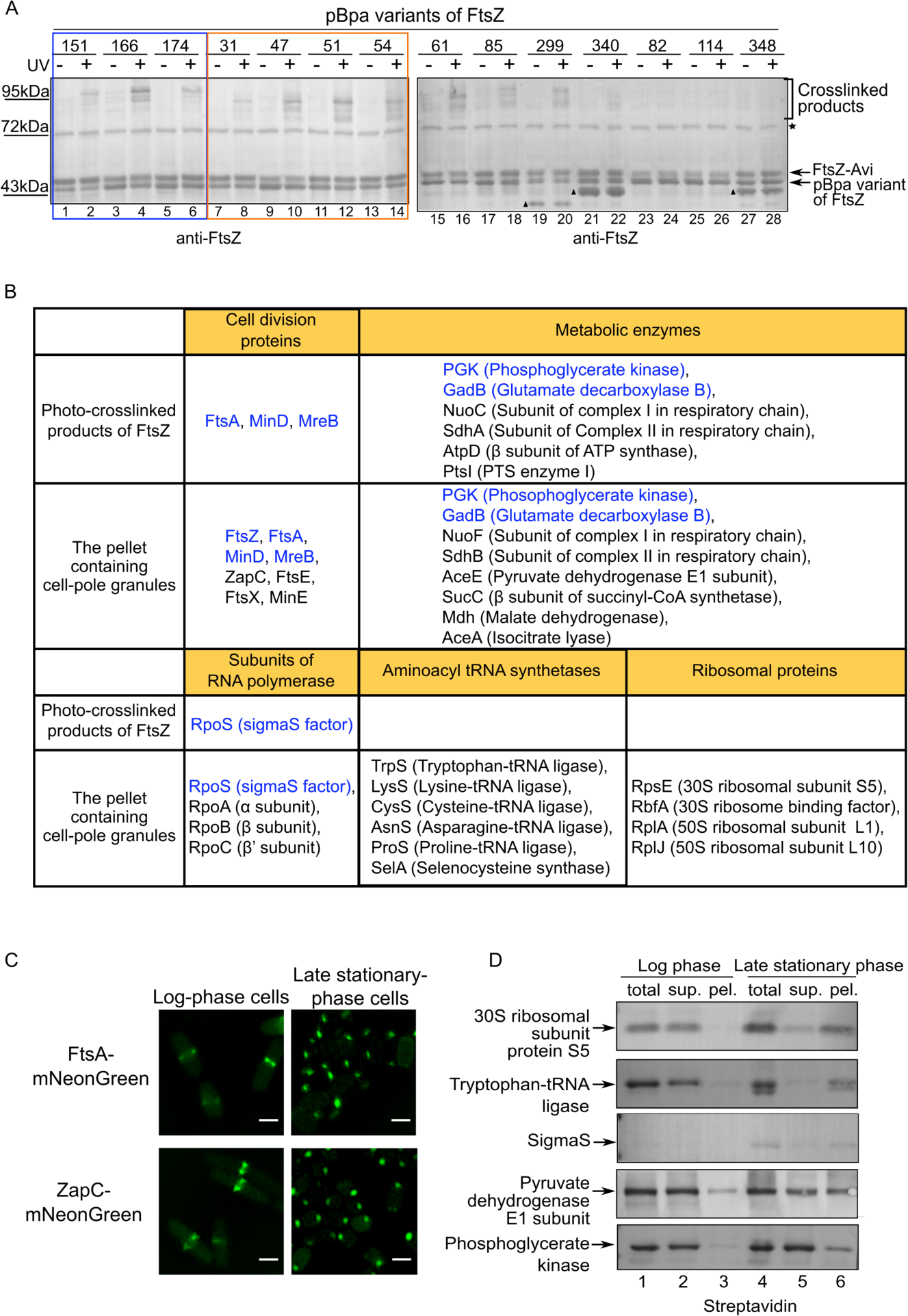
The cell-pole granules (quiescent bodies) selectively sequester proteins essential for cell growth and division. Immunoblotting results for detecting the photo-crosslinked products of the indicated pBpa variants of FtsZ in late stationary-phase *ftsZ*-*Avi* cells, as probed with antibodies against FtsZ. The asterisk indicates a non-specific protein band, and the triangles indicate the truncated forms of FtsZ produced due to the failure of pBpa incorporation. List of major proteins that were identified by mass spectrometry analysis from the photo-crosslinked products of pBpa variants of FtsZ in the late stationary-phase cells and the quiescent body-containing pellet. Fluorescence microscopic images of the log-phase or late stationary-phase *E. coli* cells expressing FtsA-mNeonGreen and ZapC-mNeonGreen. Scale bars, 1 m. Blotting results for analyzing the distribution of the indicated proteins (linked with an Avi tag for efficient detection) in the indicated fractions of log-phase or late stationary-phase wild-type *E. coli* cells, probed with streptavidin against the Avi tag. Of note, the sigmaS factor was previously reported to accumulate only in late stationary-phase but not log-phase cells (Zhou and Gottesman, 1998).

Whereas those variants having pBpa at residue positions spatially distant from each other exhibit distinct patterns of photo-crosslinked products (e.g., residues 151 and 31, or 61 and 299). In addition, no photo-crosslinked products were detected when pBpa at other residue positions in FtsZ (e.g., 82, 114 or 348) and the photo-crosslinked dimers were undetectable in these pBpa variants of FtsZ (Fig. S3A).

To identify the proteins present in the cell-pole granules, we first purified the photo-crosslinked products for each of five of these pBpa variants (with pBpa introduced at position 47, 51, 61, 140, or 166 in FtsZ) in late stationary-phase cells, via affinity chromatography, and identified their crosslinked interacting proteins by mass spectrometry analysis. Those that have a high match score and proper molecular size are listed in Fig. 2B. Interestingly, all these proteins are functionally important ones involved in cell division, metabolism and transcription.

Given that the cell-pole granules could exist as entities in the pellet fraction of the late stationary-phase cell lysates (as shown in Fig. S2C), we isolated the pellet fraction enriched with the cell-pole granules from lysed wild-type cells and identified the components present in them by mass spectrometry analysis. As also shown in Fig. 2B, all the identified proteins, some of which were also identified in the photo-crosslinked products (colored in blue in Fig. 2B), seem to be functionally important and involved in such key processes as cell division, metabolism, transcription and translation.

We subsequently verified by live-cell imaging analysis that FtsA and ZapC are indeed present in the cell-pole granules in late stationary-phase cells (Fig. 2C). In addition, we demonstrated that FtsA co-localized with FtsZ-mNeonGreen, not only in the Z-ring structure in log-phase cells but also in the cell-pole granules in late stationary-phase cells or in their lysates, as shown in Fig. S3B. In contrast to FtsA or ZapC, FtsL and ZapA, neither were identified in the cell-pole granules, were not present in the cell-pole granules (Fig. S3C). We also analyzed the distribution of several other identified proteins (that are not involved in Z-ring structure and thus difficult to be visualized by live-cell imaging) and verified their presence in the pellet of late stationary-phase cell lysates, when fused to an Avi-tag (Fig. 2D). It is worth noting that at least for ribosomal subunit protein S5 and tryptophan-tRNA ligase, both being essential for cell growth, the proteins were hardly detected in the pellet of log-phase cell lysate, but largely detected in the pellet of late stationary-phase cell lysate (lanes 3 and 6, respectively) (Fig. 1D).

Taken together, these observations suggested that the cell-pole granules seem to selectively sequester some of the proteins essential for cell growth and division in the non-growing cells. In light of this, we hereafter designate the cell-pole granules as “quiescent bodies” and continue to explore their biological significance.

In terms of their biological significance, one of the most immediate and important question is to unveil the advantages quiescent bodies would provide for the bacterial cells. The difficulty is that the dozens of proteins we identified above in the quiescent bodies are mostly not stationary-phase specific and, further, they are involved in multiple biological processes. This prevented us to put out any plausible hypothesis on their biological significance. We then decided to first find normal or mutant bacterial variants that possess various degree of quiescent body formation and then to examine possible differences in their properties.

### The formation of quiescent bodies occurs in a progressive manner in each cell and in a heterogeneous manner among the individual isogenic stationary-phase cells, and can be effectively promoted in the presence of indole but dramatically reduced upon a knockdown of the respiratory chain gene *nuoA* or *sdhC*

As a start, we tried to track the time-dependent formation of quiescent bodies in the *ftsZ*-*mNeonGreen* cells that were cultured to particular time points from log-phase to late-stationary phase in the presence of rhamnose (to induce the production of FtsZ-mNeonGreen), again by performing live-cell imaging analysis. The data displayed in Fig. 3A revealed, remarkably, that the formation of quiescent bodies appears to be highly progressive in each single cell and highly heterogeneous among individual cells. Specifically, at the 12 h culturing point, the Z-ring became undetectable in many cells and the quiescent body was not yet visible in them. When cultured for 15 h, the quiescent body with small size could be visualized in some cells lacking the Z-ring (indicated by the white arrow), while the Z-ring was still visible in a small part of others (indicated by the red arrow). From 18 h culturing to 24 h, the quiescent body was formed in more cells and the size of quiescent body became larger gradually.

**Figure 3.**
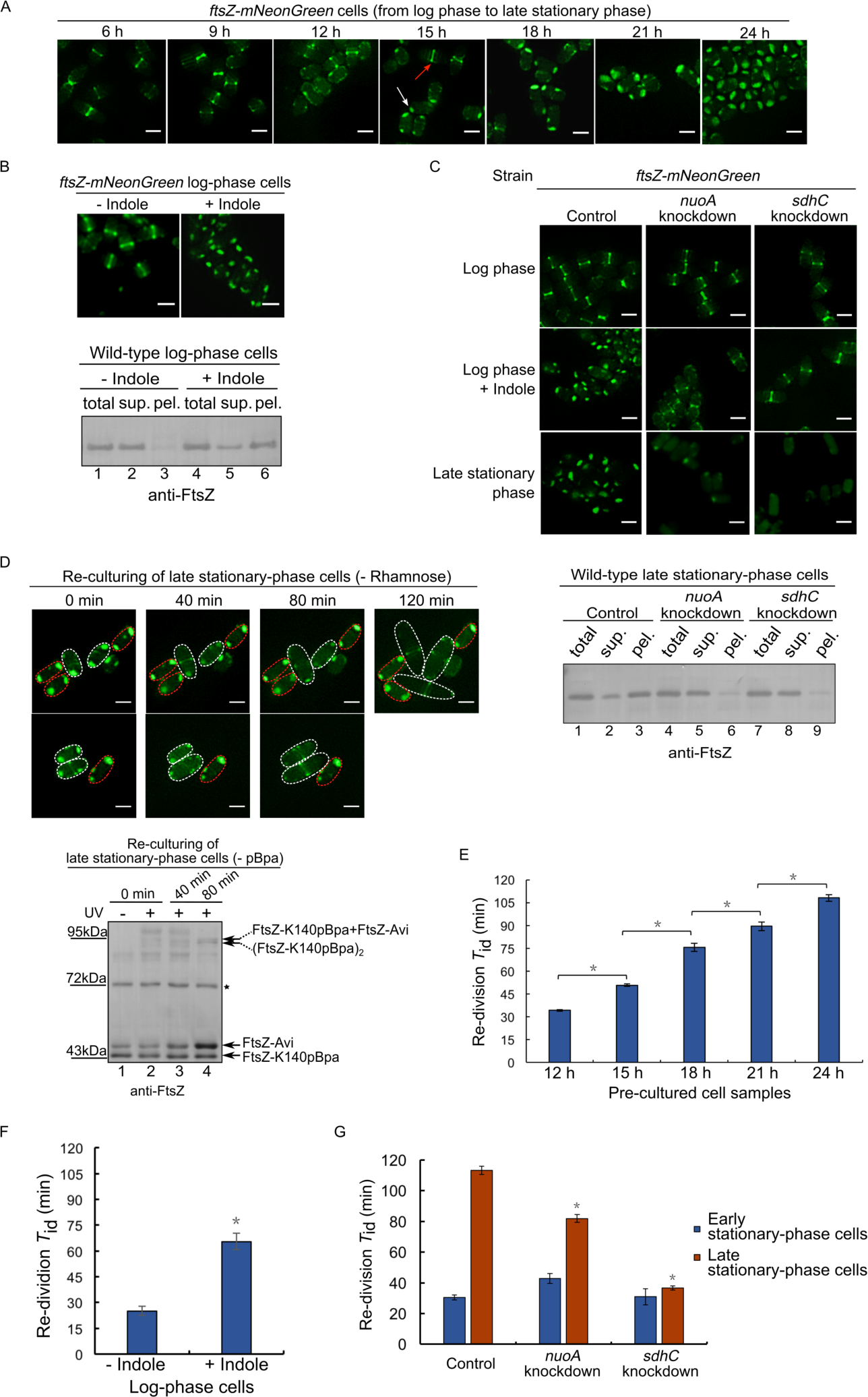
The FtsZ sequestered in the quiescent bodies is reutilized and the degree of quiescent body formation is strongly correlated to duration of the lag time upon the re-growth of bacterial cells. Fluorescence microscopic images of live *ftsZ-mNeonGreen* cells as recorded at the indicated culturing points. The bacterial cells were grown in Luria-Bertani (LB) culture medium containing 0.02% rhamnose (to induce the production of FtsZ-mNeonGreen). Scale bars, 1 m. (top panel) Fluorescence microscopic images of live log-phase *ftsZ-mNeonGreen* cells that were treated with indole at 5 mM for 1 hour. As negative control (- indole), the cells were treated with DMSO (the solvent used for dissolving indole). Scale bars, 1 m. (bottom panel) Immunoblotting results for detecting the presence of FtsZ in the total cell lysate (total), supernatant (sup.) or pellet (pel.) fraction of the log-phase wild-type cells that were treated with indole, as probed with antibodies against FtsZ. (top panel) Fluorescence microscopic images of live log-phase (indole treated) or late stationary-phase *ftsZ-mNeonGreen* cells having a knockdown of either the *nuoA* or *sdhC* gene. For the control, a non-targeting crRNA (CRISPR RNA) was expressed in the cells. Scale bars, 1 m. (bottom panel) Immunoblotting results for detecting the distribution of FtsZ in the indicated fractions of late stationary-phase *nuoA*- or *sdhC*-knockdown, or wild type (control) cells, as probed with anti-FtsZ antibodies. (top panel) Fluorescence microscopic images of live late stationary-phase *ftsZ-mNeonGreen* cells that were re-cultured to the indicated time points in fresh LB medium lacking rhamnose. (Note: one of the re-growing cells in the upper row divided into two daughter cells at the 120 min re-culturing time point). Scale bars, 1 m. (bottom panel) Immunoblotting results for detecting photo-crosslinked products of FtsZ-K140pBpa in late stationary-phase *ftsZ-Avi* cells after being re-cultured to the indicated time points in fresh pBpa-lacking LB medium, as probed with antibodies against FtsZ. The re-division *T*_id_ values of wild-type stationary-phase cells that were pre-cultured to the indicated time points in stationary phase. The cells were re-cultured (after diluting 40-fold) at 37^o^C in fresh LB medium. The re-division *T*_id_ values were calculated based on the increase in cell number within the first 30 min of the re-culturing (for details, see Methods). The re-division *T*_id_ values of wild-type log-phase cells that were untreated or treated with indole (5 mM, 1 h). The re-division *T*_id_ values of early (blue bars) or late (red bars) stationary-phase wild-type control (in which a non-targeting crRNA was expressed from a plasmid), the *nuoA*- or *sdhC*-knockdown cells.

In view that the quiescent bodies were formed in the bacterial cells only after certain culturing duration, we supposed that the late stationary-phase culture medium might be able to induce their formation in log-phase cells. This hypothesis was somehow proved by our data shown in Fig. S4A, which indicate that formation of quiescent bodies in the actively growing log-phase cells could be effectively promoted by the late stationary-phase culture medium. We then demonstrated that indole, as a specific metabolic product accumulates in the late stationary-phase culture medium (Sezonov et al., 2007), was also able to effectively promote the formation of quiescent bodies in the log-phase cells (Fig. 3B, top panel). In line with this accelerated formation of quiescent bodies, a large portion of the FtsZ protein was detected in the pellet fraction of such indole-treated log-phase wild-type cell lysate (Fig. 3B, bottom panel).

During the indole induction experiment, we accidentally noticed that quiescent bodies would not form if the log-phase cells were kept in airtight tubes without shaking, i.e., with a lack of adequate oxygen, in the presence of indole (Fig. S4B). Given that indole has been reported to dissipate the proton gradient across the inner membrane in *E. coli* cells (Chimerel et al., 2013), such an inducing effect of indole might be attributed to its acceleration of cellular respiration, which would explain why an adequate supply of oxygen is pivotal during indole induction. In light of this, we then tired to delineate whether quiescent body formation was affected in the *ftsZ-mNeonGreen* strain having a knockdown of either the *nuoA* (encodes one subunit of complex I in the respiratory chain) or *sdhC* (encodes one subunit of complex II) gene as brought about via the CRISPRi technology (Luo et al., 2015). The live-cell imaging results, shown in Fig. 3C (up panel), revealed that formation of quiescent bodies only scarcely occurred in the *sdhC*-knockdown or rarely in the *nuoA*-knockdown late-stationary phase cells. Consistently, formation of quiescent bodies no longer occurred in the log-phase cells of either the *sdhC-* or *nuoA*-knockdown strains when treated with indole. Of note, we obtained similar results with the *nouAB*-or *sdhCDAB-*knockout *E. coli* mutant strains (Fig. S4C). In line with these failures of quiescent body formation, the immunoblotting data showed little endogenous FtsZ protein in the pellet fraction of the wild-type late stationary-phase cells having a knockdown of either the *nuoA* or *sdhC* gene (Fig. 3C, bottom panel).

### Quiescent bodies are dissolved and the FtsZ protein is re-utilized to form the Z-ring structure when non-growing late stationary-phase cells resume growth in fresh culture medium

We subsequently tried to elucidate the biological significance of the quiescent body. First, we addressed whether the formation of quiescent bodies would provide an advantage such that it endows a higher capacity for the late stationary-phase wild-type cells to resist stress conditions in comparison with the mutant cells having a knockdown of either the *nuoA* or *sdhC* gene. Nevertheless, we found that all these cells possessed a similar high level of resistance towards such stress conditions as heat shock temperature, high osmotic pressure, and treatment with acid, SDS or antibiotics (data not shown).

In view that multiple functionally essential proteins were apparently sequestered in the quiescent bodies likely in their folded forms, we then addressed whether the formation of quiescent bodies would provide an advantage for the non-growing bacterial cells to reinitiate their growth by releasing these proteins for immediate re-functioning. Specifically, we examined the fate of the FtsZ-mNeonGreen protein stored in the quiescent bodies when the non-growing late stationary-phase *ftsZ-mNeonGreen* cells were placed in fresh culture medium. The live-cell imaging data displayed in Fig. 3D revealed, intriguingly, that when re-cultured in fresh LB medium lacking rhamnose (no new FtsZ-mNeonGreen protein would be synthesized), a time-dependent relocation of the FtsZ-mNeonGreen protein from the quiescent bodies to the Z-ring structure indeed occurred for cells that started to re-grow (cells circled with white dashed lines). By contrast, the FtsZ-mNeonGreen remained in the quiescent bodies for cells that have not yet started their re-growth (cells circled with red dashed lines in Fig. 3D).

This remarkable reversible nature for the FtsZ proteins sequestered in the quiescent bodies was also verified by our *in vivo* photo-crosslinking data. In particular, a time-dependent re-formation of photo-crosslinked FtsZ-K140pBpa dimers, accompanied by a parallel decrease in the level of photo-crosslinked products between FtsZ-K140pBpa and other proteins, was observed when the non-growing cells were re-cultured in fresh pBpa-lacking (thus synthesis of new FtsZ-K140pBpa protein was avoided) LB medium (Fig. 3D, bottom panel, lanes 2-4). Collectively, these observations suggested that proteins (as represented by FtsZ) sequestered in the quiescent bodies could be released for re-functioning when late stationary-phase bacterial cells re-grow and re-divide.

It is noteworthy that our live-cell imaging data of *ftsZ-mNeonGreen* cells, as displayed in Fig. 3D, also demonstrated that the dissolution of quiescent bodies, analogous to their formation (as shown in Fig. 3A), similarly occurred in a highly heterogeneous manner among individual cells. Conceivably, such heterogeneities in their formation and dissolution might be related in a certain way. For example, since the formation of quiescent bodies in each stationary-phase cell apparently occurred in multiple progressive stages (Fig. 3A), it is likely that a more mature quiescent body would take longer time dissolve.

### A higher degree of quiescent body formation in the non-growing cells is correlated to a longer lag time for them to initiate the re-growth

In view of the high heterogeneity of quiescent body dissolution among individual cells upon their re-growth (Fig. 3D), we then asked whether or not the duration of the lag time for a non-growing bacterial cell population to recover is correlated to the degree of quiescent body formation in them. For this, we made use of the cells that were known to form quiescent bodies to different degrees (**Figs. 3A**, **3B** and **3C**). Besides, we decided to use the average initial doubling time upon re-division (or re-division *T*_id_ in short), which was calculated on the basis of the re-culturing growth curves displayed in **Fig. S5**, to represent the lag time.

The results displayed in Fig. 3E-3G reveal a strong correlation between the duration of lag time and the degree of quiescent body formation. First, as shown in Fig. 3E, the cell population pre-cultured for longer time in the stationary phase, which contain more mature quiescent bodies in more individual cells, display higher re-division *T*_id_ values. Second, as shown in Fig. 3F, the re-division *T*_id_ value of the indole-treated log-phase cells is about 2.5 fold of that of the non-treated log-phase cells (~65 min vs ~26 min). Third, as shown in Fig. 3G, the re-division *T*_id_ value of the *nuoA* or *sdhC* knockdown late stationary-phase cells is lower than that for the wild-type control cells (in which a non-targeting CRISPR RNA is expressed). Notably, the re-division *T*_id_ value of the *sdhC* knockdown late stationary-phase cells is largely comparable with that of its early stationary-phase cells (Fig. 3G), indicating that the non-growing cells lacking quiescent bodies would not exhibit a prominent delay during recovery when they encounter permissive conditions.

Taken together, these results strongly suggest that the degree of quiescent body formation is highly correlated with the duration of lag time, hence the quiescent body apparently functions as a biological timer for a non-growing bacterial cell to resume growth.

### A higher degree of quiescent body formation is correlated to a higher degree of antibiotic tolerance for bacterial cells

In light of the previous report that an extended lag time correlates to an increased tolerance towards antibiotics (Fridman et al., 2014), we wondered whether or not there is a correlation between the quiescent body formation and the antibiotic tolerance. As shown in Fig. 4A and Fig. 4B, the survival rates of the cells lacking quiescent bodies, e.g., non-treated log-phase, *nuoA*- or *sdhC*-knockdown late stationary-phase cells, are much lower than the cells with quiescent bodies (indole-treated log-phase or the wild-type late stationary-phase cells) when they are inoculated into fresh medium containing either the antibiotic ofloxacin or ampicillin.

**Figure 4.**
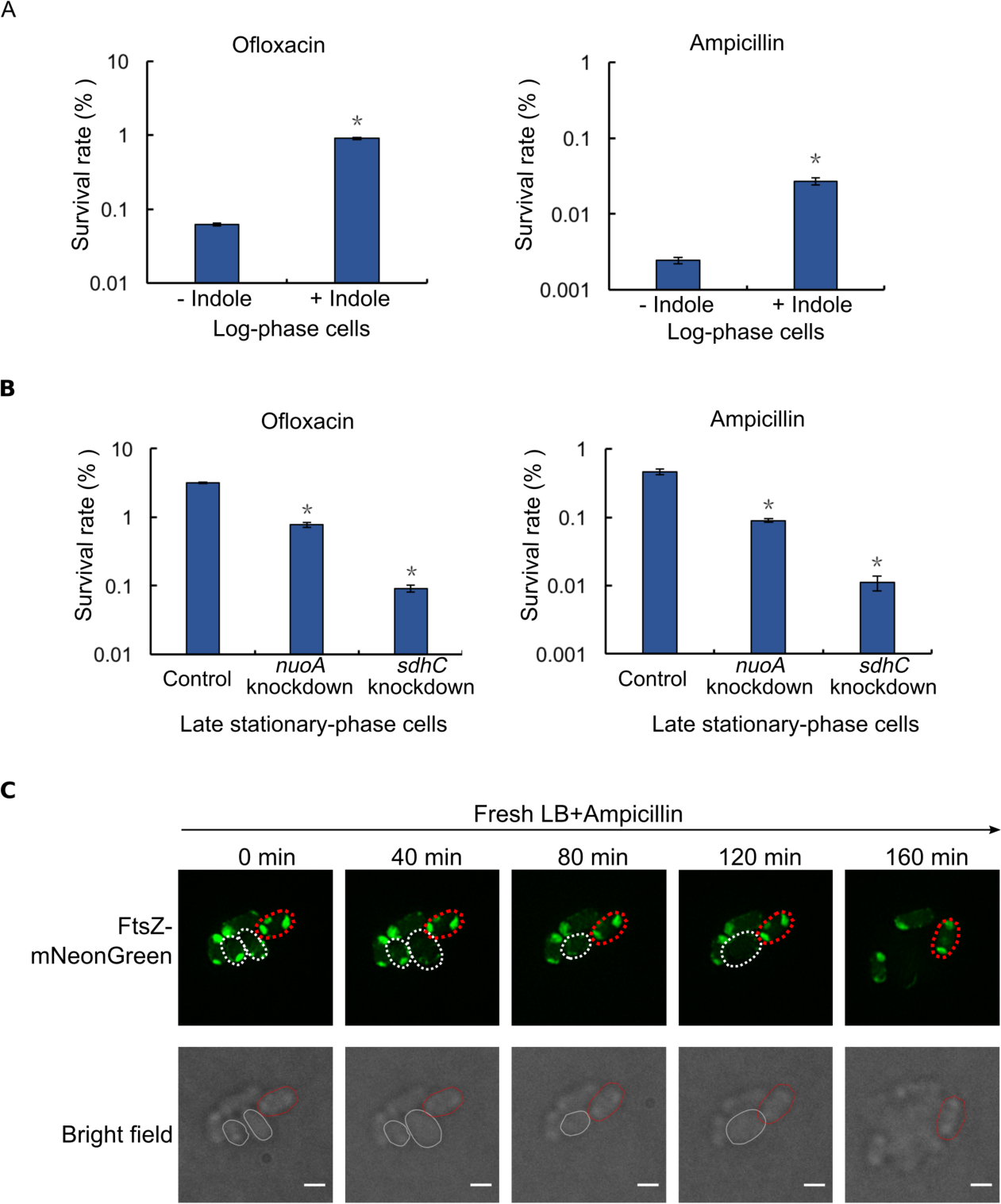
The formation of quiescent bodies is strongly correlated to the antibiotic tolerance for *E. coli* cells. Survival rates of wild-type log-phase cells that were untreated (- indole) or treated (+ indole) with indole, and subsequently re-cultured in fresh LB medium containing ofloxacin (5 μg/ml) or ampicillin (200 μg/ml). The survival rates were calculated as: [colony-forming units of the antibiotic-treated cells] / [colony-forming units of the antibiotic-untreated cells] ×100. Survival rates of the late stationary-phase wild-type (control) and *nuoA*- or *sdhC*-knockdown cells that were re-cultured in fresh LB medium containing the indicated antibiotics. Again, a non-targeting crRNA was transcribed from a plasmid in the control cells. The symbol * in all the above panels indicates a significant difference between the compared pair of samples (*P*-value <0.01, *t*-test). At least three replicates were performed for each measurement. Live-cell fluorescence (top) or bright field (bottom) microscopic images of the late stationary-phase *ftsZ-mNeonGreen* cells that were re-cultured at 37^o^C in fresh ampicillin-containing LB medium to the indicated time points. The re-growing cells and non-growing cells are marked by the white and red circles, respectively. Scale bars, 1 m.

In line with the above observations, the imaging analysis with the live *ftsZ-mNeonGreen* cells, as shown in Fig. 4C, revealed that cells (represented by the two circled with white dashed lines) whose quiescent bodies became dissolved would be eventually killed after going through a swelling process (i.e., became invisible at a certain time point after an increase in cell size) when ampicillin as present during the re-culturing process. By contrast, the cells (represented by the one circled with red dashed lines in Fig. 4C) whose quiescent bodies remained intact would survive (i.e., remained visible at their original cellular sizes at all the time points) during such re-culturing. These results implicated that the quiescent bodies would serve as an effective biomarker for identifying the rarely-exist non-growing cells in the population even in the log-phase of bacterial culturing.

### Quiescent bodies are apparently also formed in such pathogenic bacterial species as *Salmonella* Typhimurium and *Shigella flexneri*

We assume that quiescent bodies would also form in other bacterial species. As a preliminary test of this, we analyzed whether the FtsZ protein in *Salmonella* Typhimurium and *Shigella flexneri*, two pathogenic bacteria respectively causing gastroenteritis and diarrhea in humans (Graham, 2002; Jennison and Verma, 2004), exhibits a growth-dependent distribution change similar to that in *E coli* cells. Our immunoblotting results, displayed in Fig. 5A, clearly demonstrated that the FtsZ protein in both species was detected largely in the pellet fraction (lanes 6 and 12) of late stationary-phase cells, whereas hardly any detected in the pellet fraction (lanes 3 and 9) of log-phase cells. We also demonstrated the strong correlation between the degree of quiescent body formation and the duration of lag time for these pathogen bacterial cells, as shown in Fig. 5B. Consistently, we observed a strong correlation between the formation of quiescent bodies and a tolerance towards antibiotics for either *Salmonella* Typhimurium or *Shigella flexneri* (Fig. 5C). These results strongly implicate that formation of quiescent bodies is a common phenomenon in non-growing bacterial cells and is correlated to the lag time duration and antibiotic tolerance for pathogenic or nonpathogenic bacteria.

**Figure 5.**
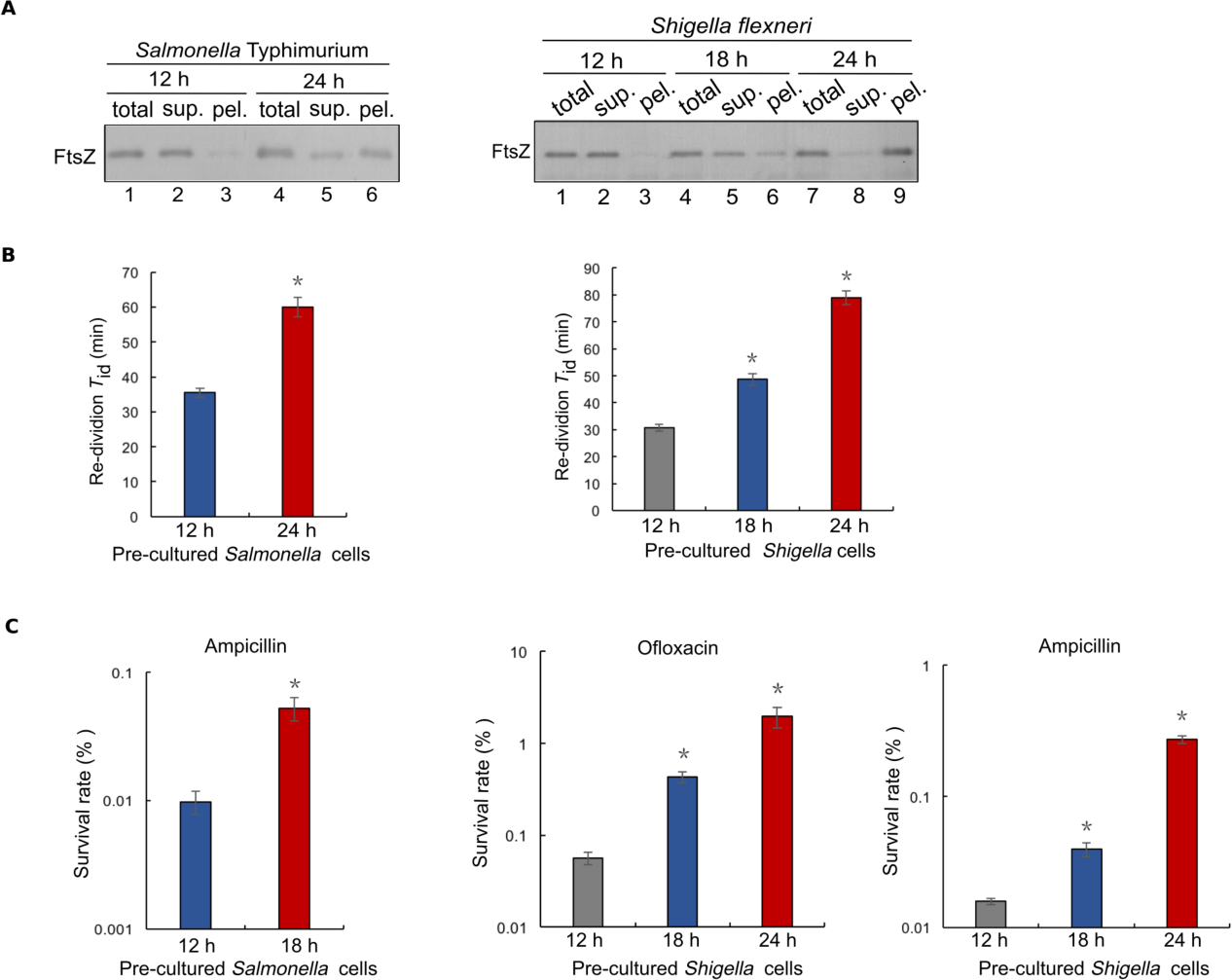
The quiescent body is apparently formed in late-stationary phase cells of pathogenic bacteria *Salmonella* Typhimurium and *Shigella flexneri*. Immunoblotting results for detecting the distribution of FtsZ in the indicated fractions of stationary-phase *Salmonella* Typhimurium or *Shigella flexneri* cells cultured to the indicated time points, as probed with anti-FtsZ antibodies. The re-division *T*_id_ values of *Salmonella* Typhimurium or *Shigella flexneri* cells that were pre-cultured to the indicated time points of stationary phase. The cells were re-cultured (after diluting 40-fold) at 37^o^C in fresh LB medium. Survival rates of the *Salmonella* Typhimurium or *Shigella flexneri* cells that were pre-cultured to the indicated time points of stationary phase and were then re-cultured in fresh LB medium containing the indicated antibiotics. The symbol * in all the above panels indicates a significant difference between the compared pair of samples (*P*-value <0.01, *t*-test). At least three replicates were performed for each measurement.

## Discussion

Here, we report the discovery of a novel reversible subcellular structure that we designated as quiescent body in bacterial cells. This structure was formed only in non-growing late stationary-phase cells and began to dissolve when the cells re-grew/re-divided in fresh culture medium. In retrospect, these findings were made as a result of our unique approaches, i.e., *in vivo* protein photo-crosslinking in combination with live-cell imaging, as well as our focus on the unique FtsZ protein, which assembles into the highly visible Z-ring structure in a dynamic and cell state-dependent manner.

One major implication of these findings is that the quiescent body apparently functions as a biological timer for a non-growing bacterial cell to resume growth, as indicated by the following observations. First, quiescent body formation among individual cells were highly heterogeneous such that it began at different culturing time points in different cells and that it apparently occurred in a progressive manner for the two quiescent bodies to form and mature in each cell (Fig. 3A). Second, the dissolution of quiescent bodies also occurred in a highly heterogeneous manner among individual cells such that they started to disassemble in some cells at a very early culturing time point while remained intact in other cells at a very late culturing point (Fig. 3D). Third, the duration of lag time for the re-culturing was strongly correlated to the degree of quiescent body formation such that the lag time is longer for cells that were derived from a later stage of the stationary phase (Fig. 3E). The role of the quiescent body as a biological timer for bacterial cell growth resumption might be reflected as such that the initiation of the dissolution would occur more efficiently and thus take less time for the “younger” quiescent bodies, whereas occur less efficiently and thus take a longer time for the “older” quiescent bodies. In other words, the differential formation and dissolution of quiescent bodies in individual cells may follow a “last-in-first-out (or first-in–last-out)” rule (Jõers and Tenson, 2016). The heterogeneous formation and dissolution of quiescent bodies in bacterial cells are schematically illustrated in Fig. 6.

**Figure 6.**
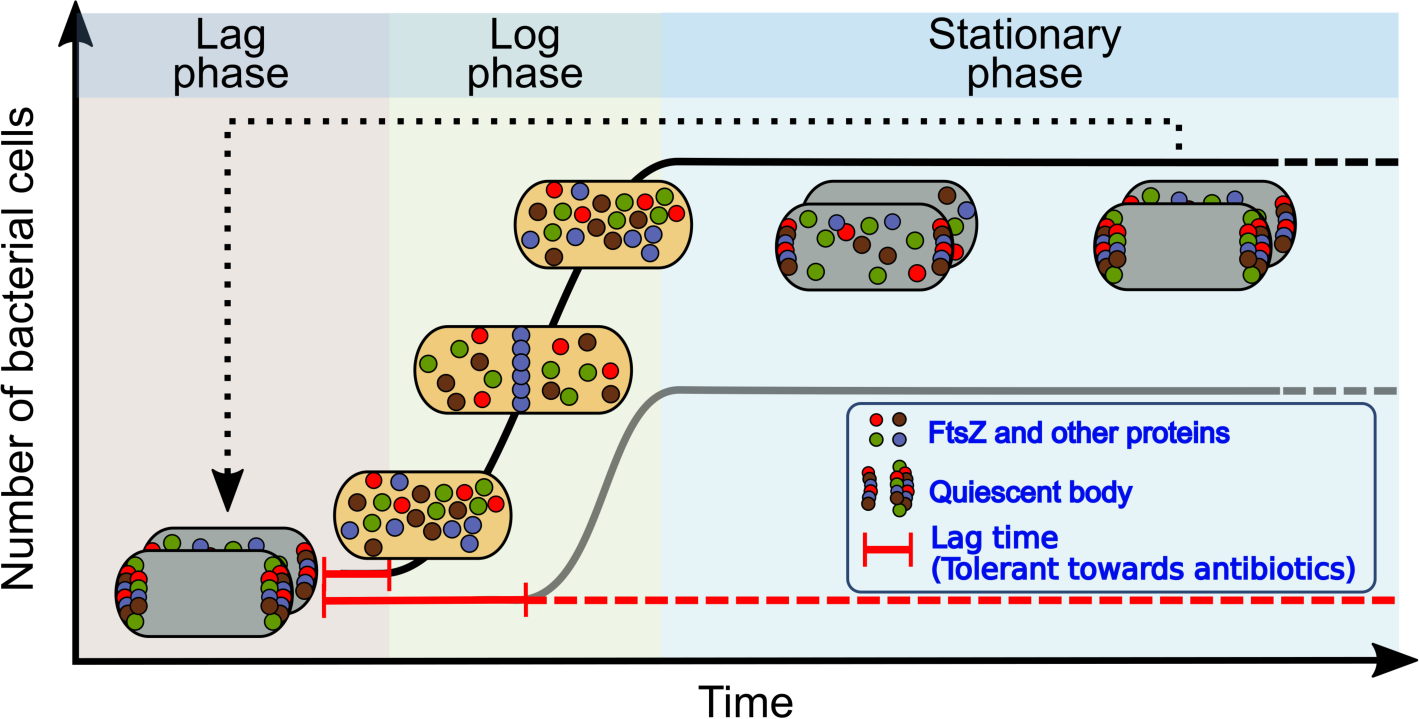
The quiescent bodies are formed in non-growing stationary phase and are dissolved in the lag-phase of bacterial cells, both in a highly heterogeneous manner. They may function as a biological timer for the non-growing bacterial cells to resume growth.

It has long been recognized that when bacteria are cultured in the laboratory, there always exists a lag phase, during which cell growth is hardly appreciable, before they resume growth from the non-growing state (Penfold, 1914). The status of the cells in this lag phase remains poorly understood, mainly due to their lack of metabolic activities. Given that a large number of proteins essential for cell growth and division are sequestered in the quiescent bodies, their formation would meanwhile lock the cells in a non-growing state. It follows that the dissolution of quiescent bodies and the release of the key proteins during the lag phase represent a major barrier for the cells to overcome before they can re-initiate their growth.

It has been known that dormant (persister) bacterial cells usually exist in an extremely low number and often possess an appearance hardly distinguishable from the non-dormant (non-persister) cells (Lewis, 2010). Owing to these obstacles, it has been greatly challenging to effectively explore this sub-population of cells and in turn to eradicate the multidrug-tolerant pathogenic persister cells. In light of our findings reported here, we propose that the formation of quiescent bodies is a far more reliable distinguishing feature for defining and identifying dormant (persister) bacterial cells, in comparison with the commonly used non-growing feature (Burke et al., 1925; Lewis, 2007). As a matter of fact, we observed in this study that wild-type cells cultured to 12 h and the *sdhC*-knockdown cells cultured to 24 h were both in a non-growing state but did not possess quiescent bodies, whereas both were able to re-grow and re-divide immediately without going through a lag phase when re-cultured in fresh LB medium (**Figs. 3E** and **3G**, respectively). This observation implicated that the non-growing feature dose not seem to make a reliable criterion for defining dormant (persister) bacterial cells. Additionally, it is commonly believed that many bacterial species collected from the natural environment exist in a “viable but non-culturable” state (Pinto et al., 2015). It is certainly worth future investigation to elucidate whether the presence of quiescent bodies is also a common and distinguishing feature of such non-growing bacterial cells.

Our data also provide some preliminary hints on how quiescent bodies are formed, as summarized below. First, the live-cell imaging data suggest that the quiescent bodies probably physically associate with the inner membrane of bacterial cells (Fig. 1E). Second, our mass spectrometry analysis identified four subunits of the inner-membrane bound complexes I and II in respiratory chain as the components of quiescent bodies (Fig. 2B). Third, a knockdown or knockout of subunits in either complex I or II resulted in a dramatic decrease of quiescent body formation (Fig. 3C). These observations implicate that the respiratory complexes on the inner membrane may play a direct role in promoting the formation of quiescent bodies. Whether the respiratory chain complexes function as a scaffold that is able to selectively recruit other proteins for facilitating the quiescent body formation is undoubtedly worth future explorations.

Much needs to be further clarified on the biology of this novel reversible subcellular structure. For example, first, whether molecules other than proteins are present in quiescent bodies? Second, how are the chemical components in the quiescent bodies organized (as might be revealed by high resolution electron microscopic studies)? Third, what are the key signal molecules that directly trigger their formation in stationary-phase cells and how are such signals sensed by the cells to initiate the molecular events leading to their formation? Fourth, how are the proteins sequestered in them specifically selected? Fifth, what signals trigger the initiation of their dissolution when the cells are exposed to growth-permissive conditions and what are the molecular events leading to their effective dissolution? Last but not least, we need to determine whether a structure similar to quiescent body is in one way or another present in single-celled or other eukaryotes.

## Author Contributions

Jiayu Yu and Yang Liu designed and performed the experiments, analyzed the data, and drafted the manuscript. Huijia Yin designed and performed the experiments. Prof. Zengyi Chang supervised this study and edited the manuscript.

## Acknowledgments

We thank Harold P. Erickson (Duke University, USA) for kindly providing us the plasmid pJSB100 and Peter G. Schultz (The Scripps Research Institute, USA) for providing us the plasmids carrying the genes encoding the orthogonal tRNA and amino acyl-tRNA for incorporating pBpa into target proteins. We thank Keio Collections for providing us the wild-type *E. coli* strain. We thank Dr. Xiaoyun Liu (Peking University, China) for generously providing us the *Salmonella* Typhimurium and *Shigella flexneri* strains. We thank the Core Facilities at the School of Life Sciences, Peking University, for assistance in using the structured illumination microscopy (SIM), and we are grateful to Drs. Chunyan Shan and Xiaochen Li for their kind help in performing the fluorescence microscopic imaging analysis. We thank Dr. Wen Zhou at the Mass Spectrometry Facility of the National Center for Protein Sciences at Peking University for assistance in performing the mass spectrometry analysis. We thank Prof. Chong Liu from Zhejiang University and Prof. Xinmiao Fu from Fujian Normal University for useful discussions. This work was supported by funds from the National Natural Science Foundation of China (No. 31670775 and 31470766 to ZYC), the National Basic Research Program of China (No. 2012CB917300 to ZYC), and the Qidong-SLS Innovation Fund. We declare that we have no conflicts of interest related to this work.

## Figure Legend

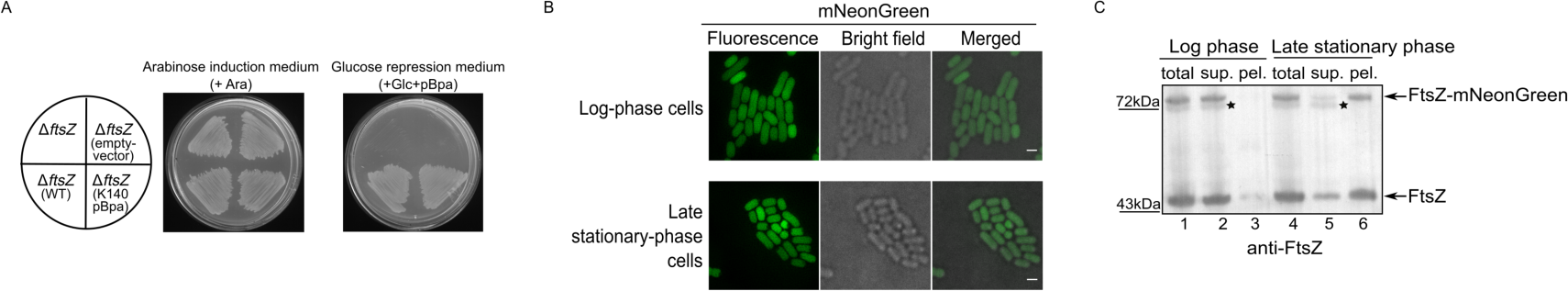

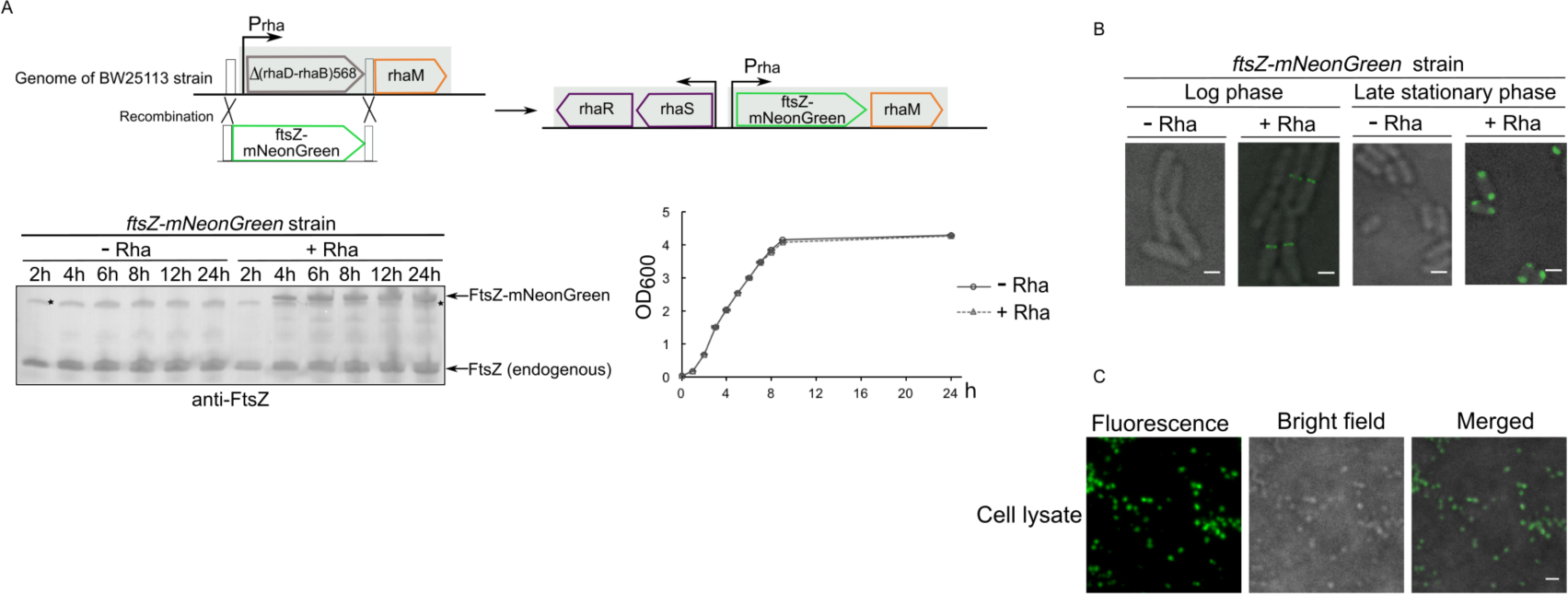

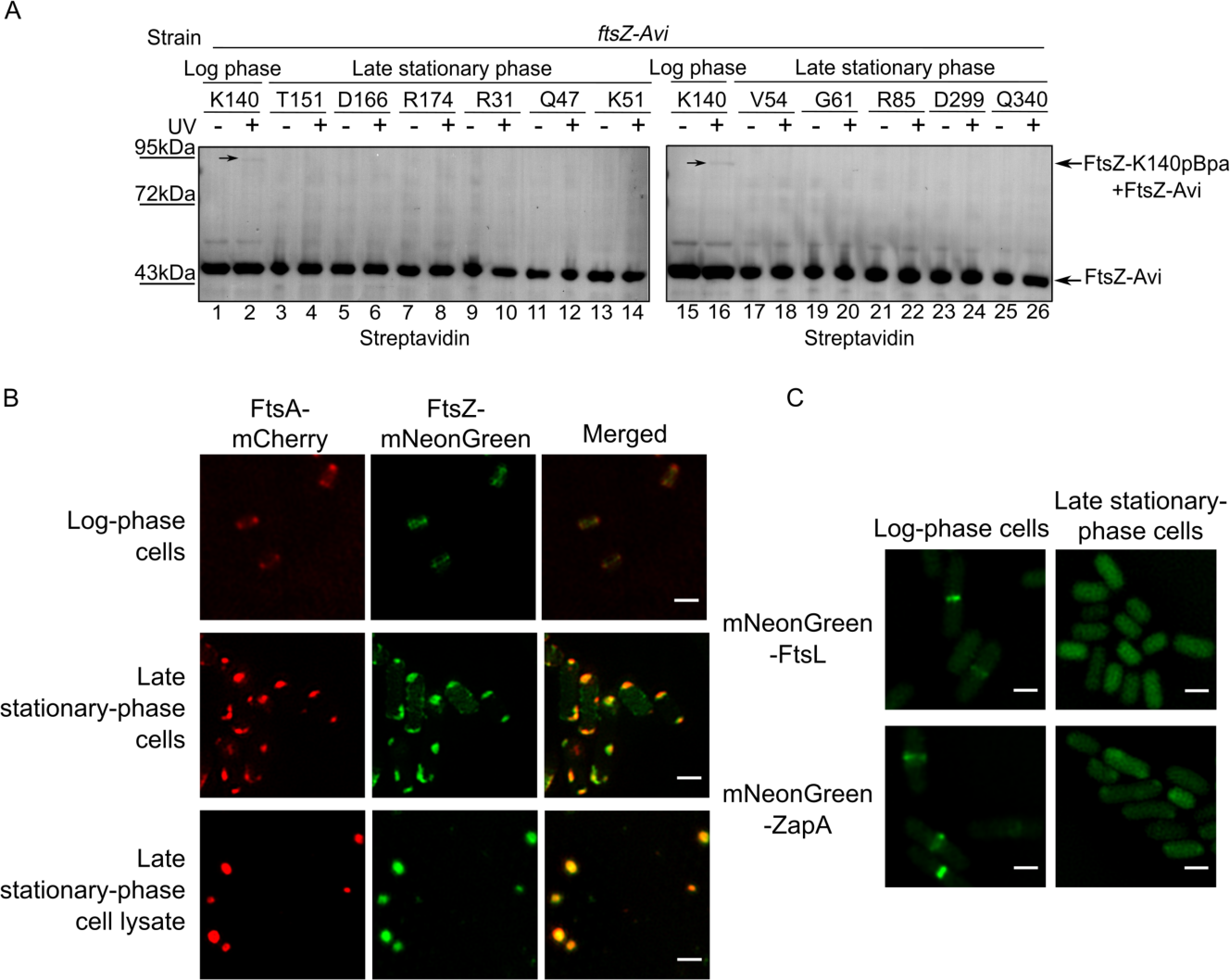

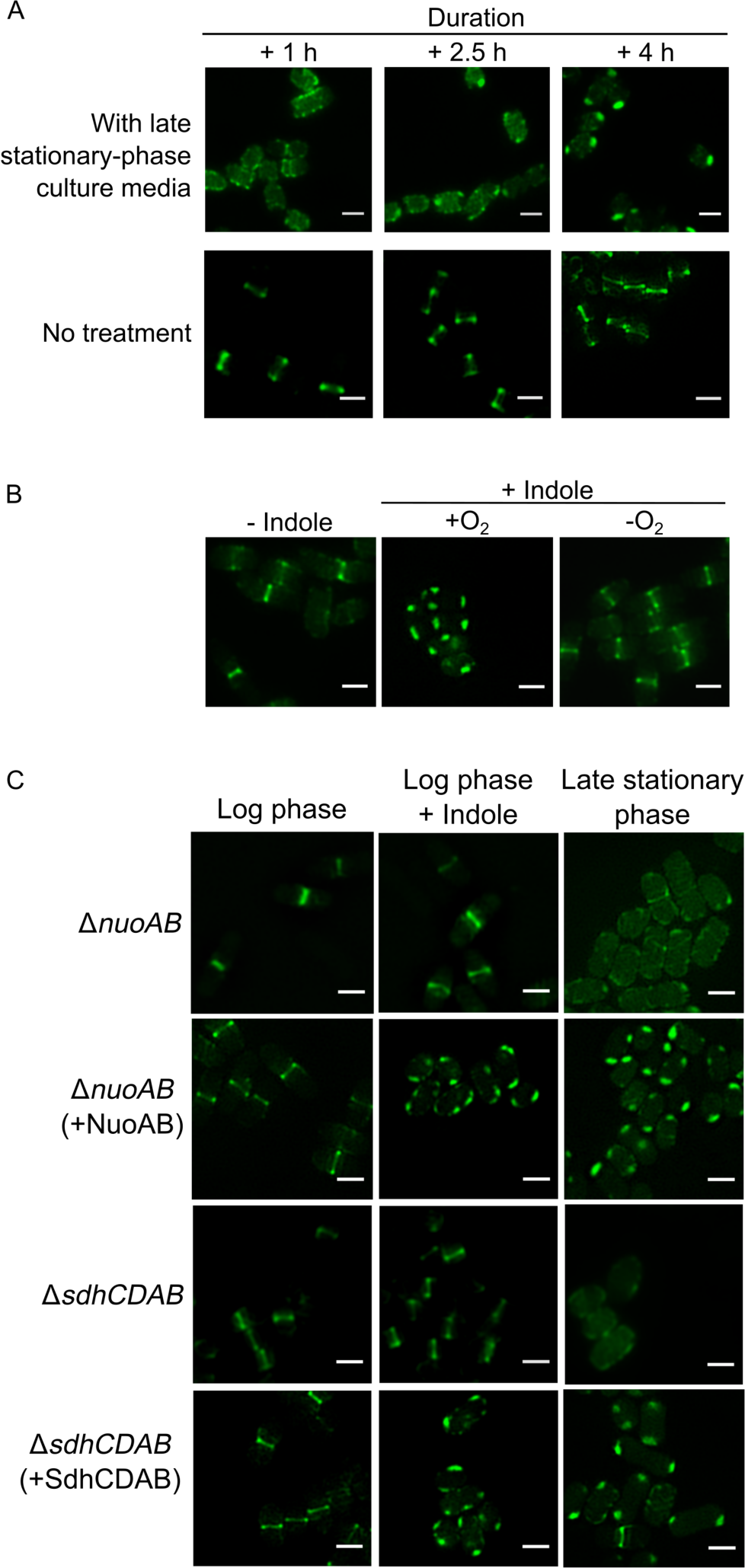

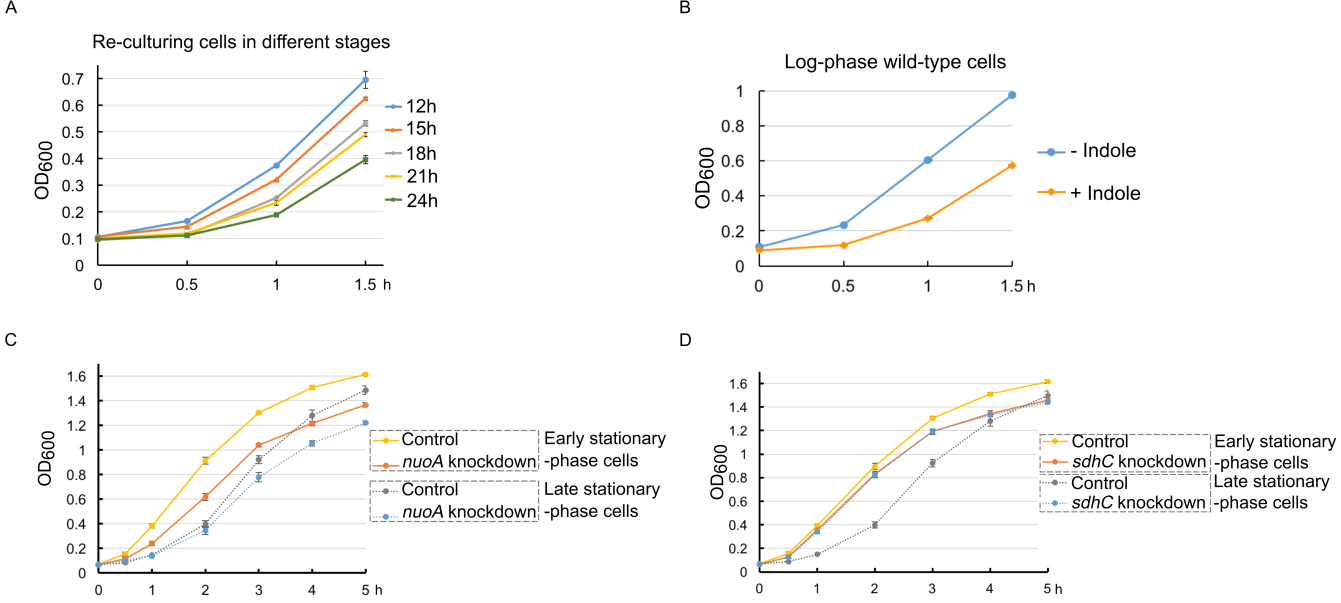

